# APPRIS principal isoforms and MANE Select transcripts in clinical variant interpretation

**DOI:** 10.1101/2021.09.17.460749

**Authors:** Fernando Pozo, Jose Manuel Rodriguez, Jesus Vazquez, Michael L. Tress

**Affiliations:** Bioinformatics Institute, Spanish National Cancer Research Centre (CNIO), Madrid, 28029, Spain; Cardiovascular Proteomics Laboratory, Centro Nacional de Investigaciones Cardiovasculares Carlos III (CNIC), 28029 Madrid, Spain; CIBER de Enfermedades Cardiovasculares (CIBERCV), 28029 Madrid, Spain

## Abstract

Most coding genes are able to generate multiple alternatively spliced transcripts. Determining which of these transcript variants produces the main protein isoform, and which of a gene’s multiple splice variants are functionally important, is crucial in comparative genomics and essential for clinical variant interpretation.

Here we show that the principal isoforms chosen by APPRIS and the MANE Select variants provide the best approximations of the main cellular protein isoforms. Principal isoforms are predicted from conservation and from protein features, and MANE transcripts are chosen from the consensus between teams of expert manual curators. APPRIS principal isoforms coincide in over 94% of coding genes with MANE Select transcripts and the two methods are particularly discriminating when they agree on the main splice variant. Where the two methods agree, the splice variants coincide with the main isoform detected in proteomics experiments in 98.2% of genes with multiple protein isoforms.

We also find that almost all ClinVar pathogenic mutations map to MANE Select or APPRIS principal isoforms. Where APPRIS and MANE agree on the main isoform, 99.93% of validated pathogenic variants map to principal rather than alternative exons. MANE Plus Clinical transcripts cover most validated pathogenic mutations in alternative coding exons. TRIFID functional importance scores are particularly useful for distinguishing clinically important alternative isoforms: the highest scoring TRIFID isoforms are more than 300 times more likely to have validated pathogenic mutations.

We find that APPRIS, MANE and TRIFID are important for determining the biological relevance of splice isoforms and should be an essential part of clinical variant interpretation.

## Introduction

Most genes produce a range of alternatively spliced transcripts. In coding genes these can theoretically produce multiple distinct protein isoforms. GENCODE v37 [1] annotates more than 64,000 distinct translations for the 13,689 coding genes that are predicted to produce multiple protein isoforms, an average of slightly more than four and a half proteins per gene.

At the protein level, all splice isoforms are not equal. Most coding genes have a main isoform [2]. The remaining gene products, the alternative isoforms, are a mixture of functionally important proteins with distinct cellular roles or distinct expression patterns [3, 4], and possibly untranslated isoforms that will have little or no cellular relevance [5, 6].

Knowing which splice variants are functionally important at the protein level is important for comparative genomics and essential for the interpretation of clinical mutations and human variation data, but the growing complexity of the human reference sets makes these analyses problematic.

Given that coding genes can have large numbers of alternatively spliced transcripts, clinical researchers are often faced with a complex choice of coding transcripts when they detect a new mutation. The American College of Medical Genetics Laboratory Quality Assurance Committee (ACMGC) recommends mapping new variants to the “most common human transcript, largest known transcript, or tissue-specific alternatively spliced transcript” [7], and referring this transcript to a reference transcript, which could be either the longest transcript or the most clinically relevant transcript [8] in RefSeq [9]. However, they also suggest that laboratories should not limit themselves to the longest transcript, but should also evaluate the impact of mutations on alternate transcripts or extended untranslated regions [8].

Historically researchers have used the longest transcript or isoform as the reference variant. Unfortunately, the longest transcript often does not produce the most biologically relevant isoform. One example is the gene *TAFAZZIN*, named after a jockstrap-wearing Italian comic character [10]. The gene product, tafazzin, is an acyltransferase that transfers fatty acids from phospholipids to lysophospholipids [11] thereby regulating cardiolipin in the mitochondrial membrane [12], Mutations affecting tafazzin cause Barth Syndrome [10], a disorder that leads to cardiomyopathy and skeletal muscle weakness, among other outcomes [13].

*TAFAZZIN* (the gene was previously known as *TAZ*, the name was presumably changed in order to avoid confusion with the ferocious, carnivorous cartoon character of the same name) is annotated with a large number of splice isoforms. UniProtKB [14] almost always represents a gene with the longest isoform, in order to map as many features as possible. When tafazzin was added in 1997, it was represented by the sequence of the longest isoform known at the time, a protein of 292 amino acid residues known as the “full-length” isoform. The expansion of the human gene set means that the “full-length isoform” is no longer the splice variant with the largest gene product; the current version of RefSeq annotates three transcripts that would produce even longer splice isoforms [Figure 1].

**Figure 1.**
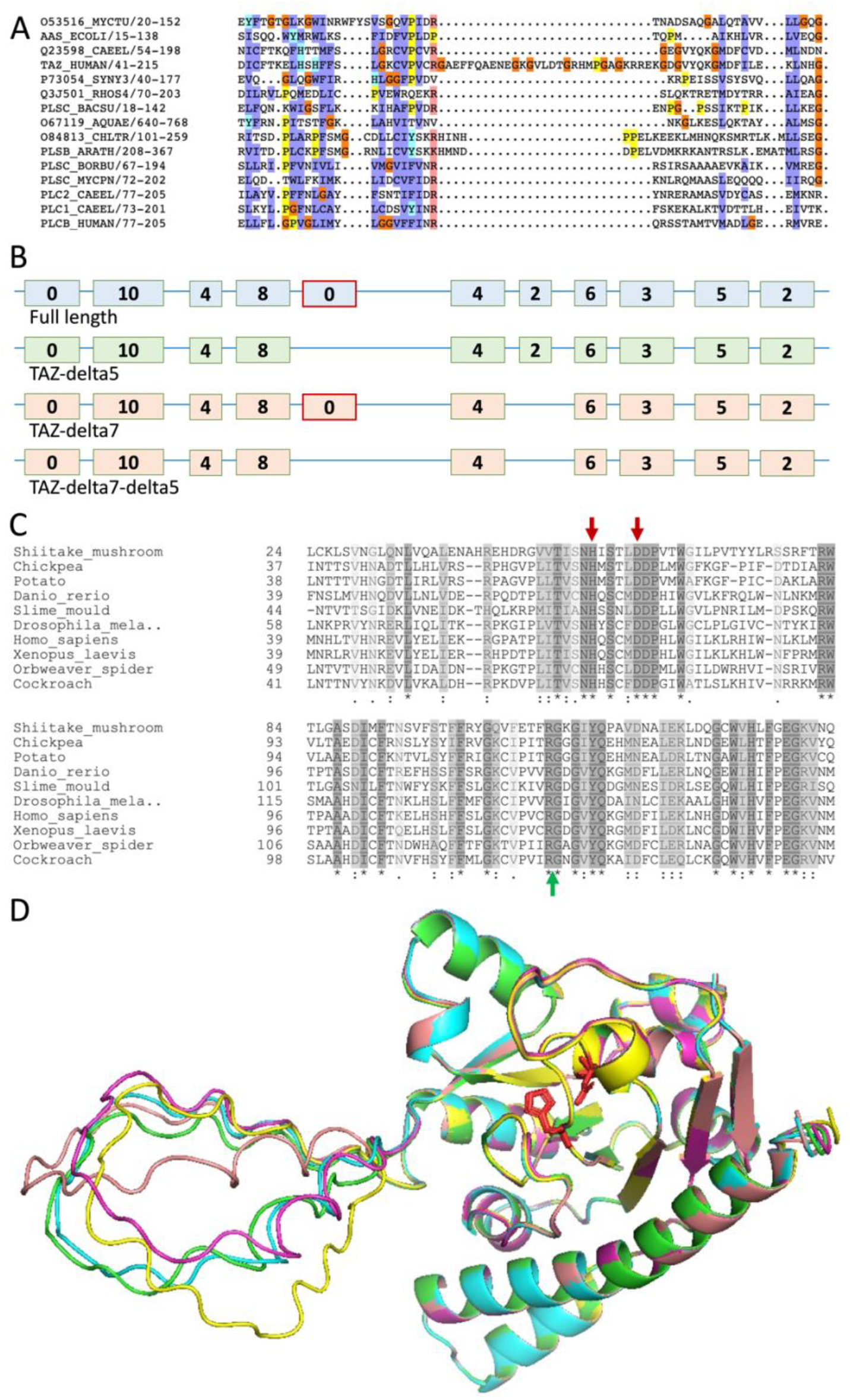
The “full-length” splice variant of *TAFAZZIN*. A. Pfam alignment showing the inserted 30 amino acids of the “full-length” variant in a section of the wider acyl transferase domain. Colours indicate residue type. B. The four most studied transcript variants of TAFAZZIN. The 11 exons of the “full-length” transcript are shown with a green background, exons from the principal transcript (TAZ-delta5) are shown with a green background. Exon 5 is highlighted in red. The number of ClinVar [18] pathogenic and likely pathogenic mutations that map to each exon is shown inside the exon. Only exon 1 (the trans-membrane helix) and exon 5 have no pathogenic mutations in ClinVar. C. The alignment of sequences equivalent to exons 2, 3, 4, 6 and 7 of the human gene from ten plant, fungi, insect and vertebrate species from UniProtKB. The probable catalytic histidine and aspartate residues are indicated by the red arrows, the insertion point by a green arrow. Conservation is indicated below the alignment (asterisked columns are entirely conserved) and by the colour of the columns, the darker the colour, the greater the conservation. D. Five models of the “full-length” isoform of *TAFAZZIN* generated by AlphaFold [19]. The models have 5 different colours and are superimposed. Likely catalytic residues are shown in stick form and coloured in red. The 30-residue insertion (on the left) is modelled as a floating loop.

In 2003, Vaz *et al* [15] produced an analysis succinctly titled “Only one splice variant of the human TAZ gene encodes a functional protein with a role in cardiolipin metabolism”. It was not the “full-length” isoform. The authors found that the functional tafazzin protein was a 262 residue protein translated from a transcript that was missing exon five of the “full-length” transcript. Further papers in 2009 [10] and 2020 [11] support this assertion. Unfortunately, UniProtKB cannot easily change the display isoform when more information becomes available. So the UniProtKB display variant remained the same.

The effects of the choice of initial main isoforms in UniProtKB can be found downstream in databases as distinct as the Pfam database [16], a database of functional protein domains, and the Locus Reference Genomic (LRG) database [17], a database of stable reference sequences designed for clinically relevant variant reporting. The Pfam database has incorporated the “full-length” isoform of tafazzin into the definition of the acyltransferase domain. The translated region from exon 5 of the “full-length” transcript stands out like a sore thumb in the 46 sequence “seed” alignment defining the domain [Figure 1], The inclusion of the exon in the acyltransferase seed alignment cements the position of the “full-length” variant as the apparent main isoform.

The *TAFAZZIN* transcript that produces the “full-length” variant was also selected as the main transcript variant in the LRG database, even though reference transcripts are chosen by expert curators. LRG did not annotate the transcript directly from UniProtKB. The reason for annotating the “full-length” transcript as the most clinically important is circular. Since early clinical researchers used the “full-length”*TAFAZZIN* transcript to report all variants, some mutations inevitably fell in exon 5. In part, LRG selects clinical representatives for genes based on capturing as many annotated mutations as possible, so it would have to choose the “full-length” transcript variant. Choosing the “full-length” transcript variant in LRG means that both the longest transcript and the most “clinically important” transcript are one and the same. And both point to a non-functional protein isoform. While 44 ClinVar variants in *TAFAZZIN* are predicted to have a pathogenic effect in Barth Syndrome, none map to exon 5 of the “full-length” transcript (Figure 1).

Conservation information also suggests that exon 5 from the full-length transcript of *TAFAZZIN* is an alternative exon. The insertion only exists in primates and the 5’ end of the exon is derived from a SINE Alu transposon. Even within the primate clade, only 9 of the 30 amino acids are conserved and the exon has poor conservation scores in both PhyloCSF [20] and PhyloP [21]. It is highly improbable that this exon has gained functional importance in such a short evolutionary time frame [22]. In addition, the insertion occurs in a region that is particularly conserved in all tafazzin proteins, across plants, fungi and animals, a region that is likely to be important in substrate binding [23]. Experiments show that the exon alters but does not remove the acyltransferase activity, although it does make the protein less stable in the mitochondrial membrane [11]. One possible effect of the insertion of 30 amino acids into the globular structure of the acyltransferase can be seen in Figure 1; the 30 amino acids in the novel loop are likely to either float freely, or stick to the surface of the protein.

That coding genes have dominant transcripts was first shown using differential transcript expression [24]. The study found that a proportion of coding genes had a transcript with five times as much expression as the next transcript across all tissues, though subsequent experiments showed that this effect was diluted with larger transcript data sets. Other methods have since sought to make use of RNA-Seq data to predict dominant splice variants [25, 26].

Most genes also have a main protein isoform at the protein level [2]. When we compared the main isoform from proteomics experiments to predicted dominant isoforms from a range of methods, we demonstrated conclusively that RNA-seq information was a poor predictor of the main cellular protein isoform [2]. Even the 5-fold dominant transcript variants from the transcriptomics experiment [24] coincided less than 80% of the time with the main proteomics isoform, while the two-fold dominant transcripts fared even more poorly. The best predictors of main cellular isoforms were unique CCDS transcript variants [27] and APPRIS principal isoforms [28], both of which coincided more than 96% of the time with the main proteomics isoform [2].

APPRIS principal isoforms are predicted from the presence and absence of protein features such as functional domains and 3D structure and cross-species conservation. APPRIS ignores non-coding regions. APPRIS chooses the 262 amino acid isoform as the main isoform for *TAFAZZIN*, though APPRIS has to make the decision based on cross species conservation. This is because both the 262 residue isoform and the “full-length” isoform score equally well for the presence of the Pfam acyltransferase domain because the “full-length” isoform was included in the definition of the acyltransferase domain [Figure 1].

CCDS transcripts are annotated by teams of expert curators from the Ensembl [29] and RefSeq [9] databases, and the same groups have recently joined forces to develop MANE [29]. For MANE, annotators use computational methods and manual review to select a single transcript per coding gene, the MANE Select transcript. Expression levels, evolutionary conservation, existence in UniProtKB and clinical significance are all balanced to determine functional potential of each isoform. Unfortunately, in the case of *TAFAZZIN*, the MANE Select transcript is the “full-length” transcript, because it is annotated in UniProtKB and supported by circular evidence from LRG and Pfam.

While all genes must have at least one functional isoform, it is not clear how many alternative splice isoforms play functional roles in the cell [5]. Although transcriptomics evidence suggests that most annotated coding transcripts are important, we found much less evidence for alternative splicing at the protein level than would be expected [30] – in fact, we detected just 0.37% of all possible detectable alternative peptides. Human genetic variation data backs this observation up: alternative exons are not, in general, under purifying selection pressure [5,31]. We have found that three quarters of alternative exons are primate-derived [4].

However, approximately 5% of alternative exons can be traced to a last common ancestor with fish [3] and these conserved alternative exons, at least, are likely to be functionally important. These ancient alternative exons are much more likely to have tissue-specific roles [4], to be detected at the protein level [3] and to have pathogenic mutations [3]. Based on these findings we developed TRIFID, a machine learning-based predictor that uses protein features and experimental evidence to predict the likely functional importance of alternative isoforms [32]. We have shown that TRIFID can discriminate functionally relevant alternative variants. Alternative exons from high scoring alternative variants are under selective pressure, while alternative exons from low scoring variants are generally not. The most discriminating feature in TRIFID is cross-species conservation.

In this study, we investigated the relationship between alternative splicing, protein expression and the annotation of clinical variants. We confirm the finding that coding genes have a single main splice isoform at the protein level. This main splice isoform almost always coincides with the MANE Select transcript and APPRIS principal isoform. APPRIS principal isoform and MANE Select transcripts also cover all but a handful of annotated pathogenic mutations in ClinVar. Approximately half of the pathogenic mutations that do map to alternative exons are either erroneous or have no evidence for pathogenicity. Most validated pathogenic mutations that map to alternative exons are from transcripts predicted to be functionally important by TRIFID, tagged as MANE Plus Clinical, or both.

## Methods

### Reference annotation

We downloaded the GENCODE v37 reference human gene set [1], both the standard gtf file for the coordinates of the coding sequences (CDS) and the coding sequence translations. We also downloaded the APPRIS [28] isoform annotations for the v37 gene set. APPRIS annotates a unique protein isoform as the principal isoform for each coding gene. APPRIS also provides TRIFID functional importance scores [32] for each annotated isoform. The normalised TRIFID scores for each isoform is an estimation of the relative functional importance of each isoform. MANE Select and MANE Plus Clinical transcripts are annotated in the GENCODE v37 reference set. MANE Select transcripts are representative transcripts agreed upon by Ensembl/GENCODE and RefSeq manual curators [29].

### Proteomics analysis

In order to distinguish dominant proteomics isoforms for as many genes as possible, we re-analysed five large-scale proteomics analyses [33–37]. The spectra from experiments PXD000561, PXD004452, PXD005445, PXD008333 and PXD010154 were downloaded from ProteomeXchange [38]. These experiments were carried out using a range of cell lines and tissue types. We set aside the experiments that did not use trypsin because other enzymes are less efficient.

We searched against the GENCODE v37 human reference set with read-through transcripts eliminated [39]. We mapped spectra to the GENCODE v37 gene set using Comet [40] with default parameters, allowing only fully tryptic peptides, one missed cleavage and the oxidation of methionines. We post-processed the peptide spectrum matches (PSMs) with Percolator [41], and the Percolator posterior error probability (PEP) values were used to order the PSMs. Valid PSMs had PEP values below 0.001. PEP values of 0.001 approximate to q-values of 0.0001, so are highly conservative. Moonlighting peptides, those that mapped to more than one gene, were discarded.

To reduce false positive identifications that will inevitably result from combining results from many different experiments [42] peptides identified in each of the five analyses had to be validated by PSMs from at least two of the multiple experiments that made up each proteomics analysis. All of these large-scale analyses contained replicate experiments, which meant that tissue-specific peptides were not excluded.

We determined the main proteomics isoform in the same way as we had in a previous analysis [2]; we simply counted up the PSMs that mapped to every annotated protein isoform within each gene. The protein isoform with the most PSM over the five analyses was determined to be the dominant experimental protein isoform (Figure 2).

**Figure 2.**
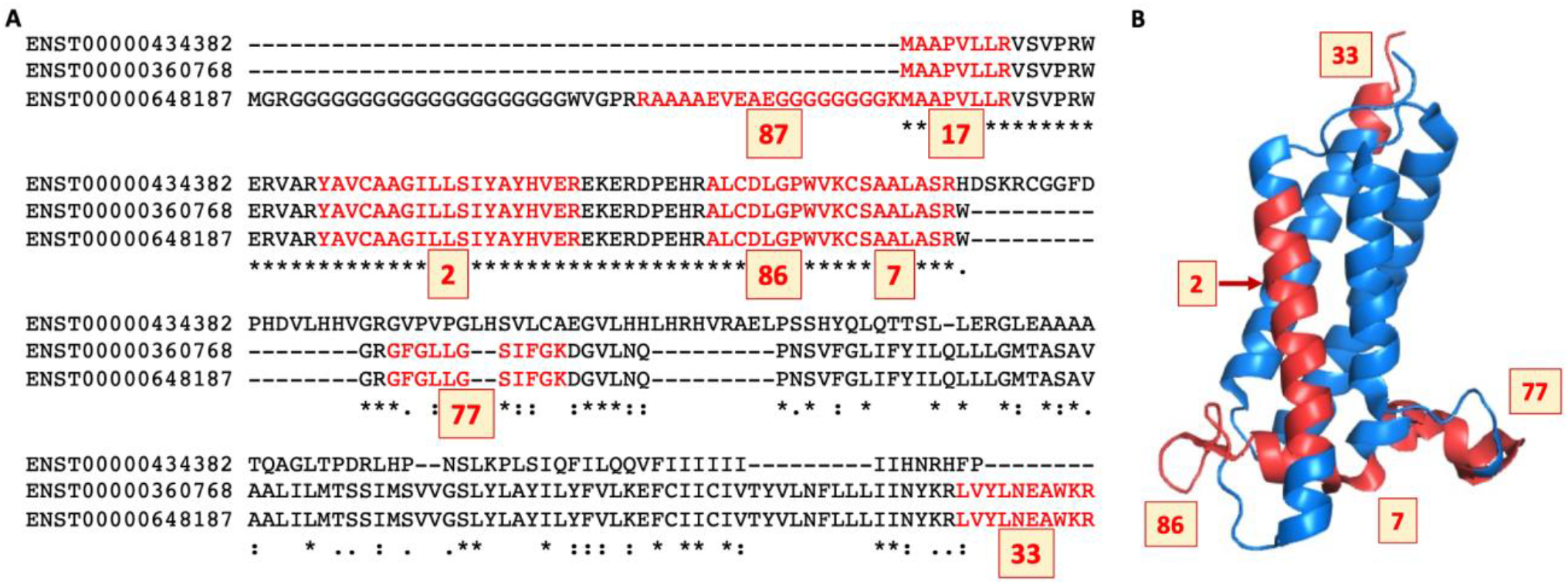
Peptides mapping to isoforms from gene *VKORC1L1*. A. The alignment between the three protein sequence unique isoforms of gene *VKORC1L1* (Vitamin K epoxide reductase complex subunit 1-like protein 1) showing the peptides that we detected in the five large-scale proteomics analyses in red. The number of PSM detected for each peptide is highlighted in the yellow boxes and the main proteomics isoform was determined by summing the peptides that mapped to each isoform. In this case, variant ENST00000648187 coded for the main proteomics isoform with a total of 309 mapped PSM. B. The peptides from *VKORC1L1* shown mapped to the structure of human *VKOR* (PDB: 6WVH, 43). Peptides are marked in red where they would fall in the real structure of *VKORC1L1* and the number of PSM that we detected are shown in blue boxes. Note that Vitamin K epoxide reductase complex subunit 1-like protein 1 is a membrane-bound protein, so except for peptide YAVCAAGIIISIYAYHVER (for which we only find 2 PSM), peptides were found exclusively for regions external to the membrane. The two N-terminal non-overlapping peptides (with 87 and 17 PSMs) do not map to the protein structure; this region is likely to be disordered. Peptide mapping to 6WVH was carried out using HHPRED [44]. The structure was generated with PyMol.

### Mapping of annotated mutations

We downloaded the April 2021 version of ClinVar and used VEP [**45**] to map the mutations to the coding genes and transcripts annotated in v37 of the GENCODE human reference set. A total of 656,237 annotated ClinVar mutations mapped to GENCODE coding variants. Along with each mutation, we recorded the official ClinVar CLIN_SIG designation, and whether or not the mutation was supported by a reference in PubMed, or was reviewed by an expert panel.

Many mutations mapped to more than one transcript. In these cases, we reduced the mapping to a single transcript for each annotated mutation. For the APPRIS analysis, when a mutation mapped to more than one transcript, we retained the mapping to the APPRIS principal transcript where possible. For the MANE analysis, we prioritized the mapping to the MANE Select transcript, and for the longest transcript analysis we prioritized the longest transcript. Mutations that did not map to a principal, MANE Select or longest transcript were associated to one of the alternative transcripts that they mapped to - in each case the alternative transcript that had the highest normalised TRIFID score.

For the analysis of pathogenic mutations, we only considered those mutations tagged as Pathogenic, Likely Pathogenic or Pathogenic/Likely Pathogenic by ClinVar. For the initial analysis we required that each of these pathogenic mutations also had a supporting PubMed publication. The reason that we insisted on PubMed references for the pathogenic mutations was to allow us to cross-check the evidence for pathogenicity for each mutation. The Richards *et al* ACMGC analysis [**8**] was excluded as a supporting PubMed publication. This paper is used as a reference for 25,382 mutations, but the paper contains no details about any of these mutations.

## Results

### Proteomics analysis

In total, we detected 337,737 distinct peptides mapping to no more than one coding gene. These peptides mapped to 15,137 coding genes and we found peptide evidence for two or more isoforms for 1,293 of these genes. Not all of these will be bona *fide* alternative splice isoforms; a proportion are likely to be caused by unfinished gene models in the GENCODE annotation.

We counted up the PSM that supported each splice isoform and chose a main proteomics isoform where possible. Main proteomics isoforms were those isoforms that were supported by at least 4 PSM and that had at least two more PSM than any other isoform annotated for the gene (see Figure 2). There were 7,697 genes where we could determine a main cellular proteomics isoform. We analysed the agreement between these main cellular isoforms and other means of selecting a main variant. We first ran a control experiment in which we selected variants randomly from each of the 7,697 genes and counted how many times they were in agreement with the main isoform. After 100 simulations, we found that random selection of isoforms agreed with the main isoform on average just 29.97% of the time (Figure 3).

**Figure 3.**
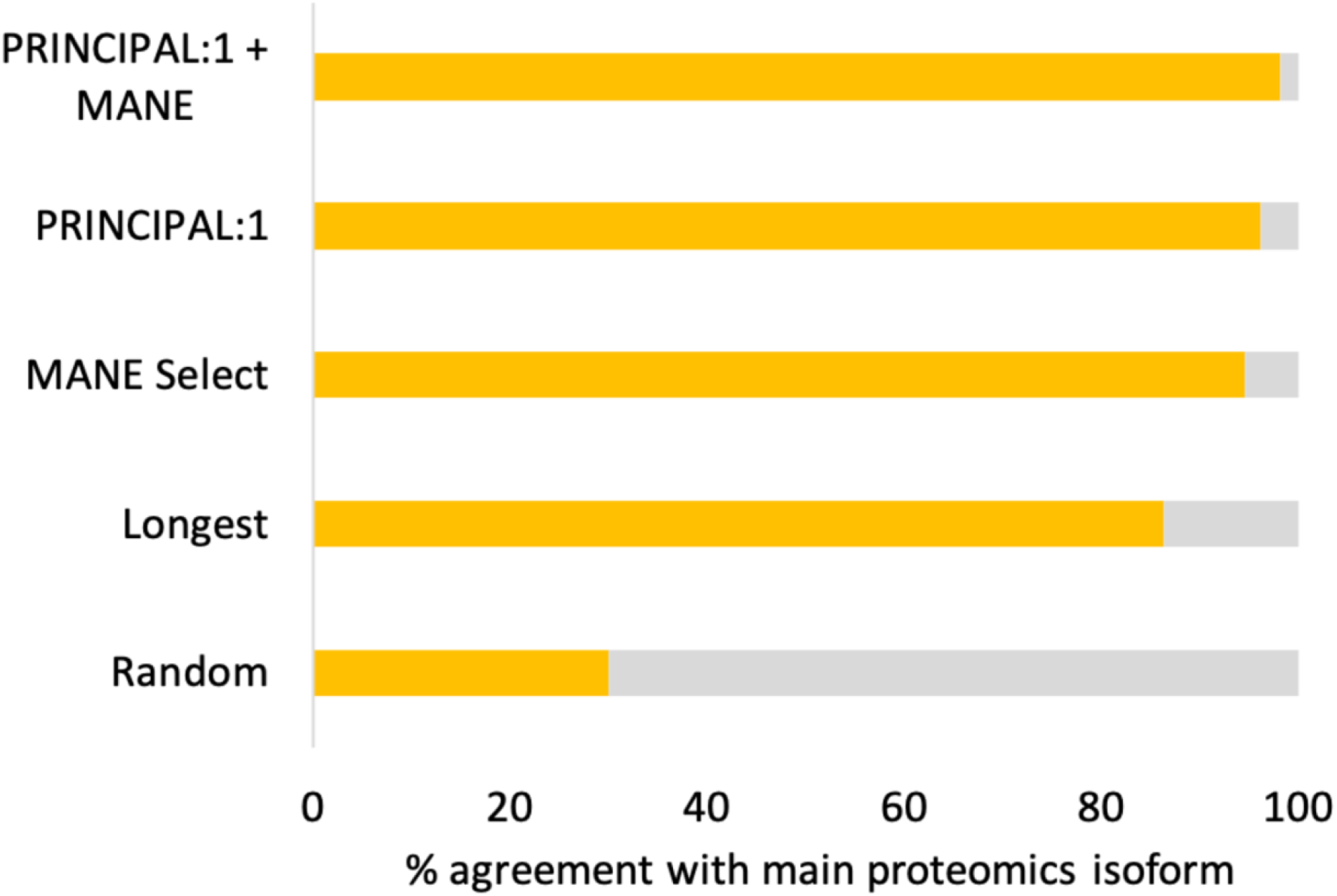
Agreement between main protein isoforms and APPRIS, longest and MANE Select variants. The percentage of genes in which predicted main isoforms coincided with proteomics main isoforms. Predicted main isoforms were the longest isoform (used by UniProtKB), the MANE Select variants (used in Ensembl/GENCODE and RefSeq gene sets) and APPRIS principal variants (available for all three databases).

Next, we compared the longest isoforms and main isoforms. The agreement between main proteomics isoforms and longest isoforms was 86.4%. The longest isoforms do capture more protein features, which is why they are used by UniProtKB [14] as the canonical isoform, but there is no biological justification for choosing the longest isoform. In this analysis the longest isoforms have an inbuilt advantage - the longer an isoform, the more peptides that can map to the isoform. This is clear in the example in Figure 2. In *VKORC1L1*, the third (longest) isoform would be produced from an upstream ATG. The first and third isoforms can be the main proteomics isoform because they have a unique sequence, so unique peptides can map to them. The second isoform (translated from ENST00000360768) cannot be the main proteomics isoform because its sequence is identical (but shorter) to that of the longest isoform. Incidentally, this second isoform is the one selected by APPRIS and MANE for this gene.

Two methods of selecting the main variant for coding genes that do have greater biological justification are APPRIS [28] and MANE. Both methods are incorporated within the Ensembl/GENCODE reference human gene set. APPRIS selects principal isoforms based on conservation of protein features and cross-species conservation [45]. MANE Select variants are generated from the agreement between the manually annotated gene models in the RefSeq and Ensembl/GENCODE reference gene sets [29].

The agreement between the MANE Select variants and the main proteomics isoform was 94.58% (Figure 3) over the 6,995 genes with MANE Select variants (as of GENCODE v37, not all genes had MANE Select variants). Meanwhile, the agreement between APPRIS P1 principal isoforms (those chosen by the core APPRIS core methods) was 96.13% over the 6,285 genes that had P1 isoforms (Figure 3). P2 isoforms are chosen using the TRIFID functional isoform predictor [31], which has recently been incorporated into APPRIS. The agreement between P1 and P2 isoforms and the main cellular isoform was 95.52% over 6,847 genes. APPRIS P3 isoforms are chosen based on proteomics evidence, so cannot be compared.

In those 5,888 genes in which the APPRIS P1 isoform and MANE Select isoform coincided, the agreement with the isoform with most proteomics evidence was 98.2%. Variants where the APPRIS P1 isoform and the MANE Select transcript agree are very good predictors of the main cellular isoform.

### ClinVar annotations

There are fewer clinical mutations annotated in alternative exons than might be expected by chance. One in 40 bases (2.43%) are exclusive to alternative exons (as defined from APPRIS), so we might expect that approximately 2.5% of all mutations would map to alternative exons by chance. In fact, the proportion is slightly smaller, 1.6% of all ClinVar mutations map to coding exons defined as alternative by APPRIS.

The breakdown by clinical significance paints a different picture. When we divided the mutations in alternative exons by their official ClinVar clinical significance designation, we found that the more damaging the class of mutations, the fewer mutations that mapped to APPRIS alternative exons. For example, just over 2.5% of mutations tagged as “Benign” map to alternative exons, close to what would be expected if mutations were evenly distributed between principal and exons (Figure 4). The percentage drops among variants of unknown significance (VUS, 1.56%) and is even lower among those mutations officially tagged as “Pathogenic” (0.89% of pathogenic mutations mapped to alternative exons). Mutations reviewed by an expert panel are least likely of all to map to alternative variants; just 0.31% map to APPRIS alternative exons.

**Figure 4.**
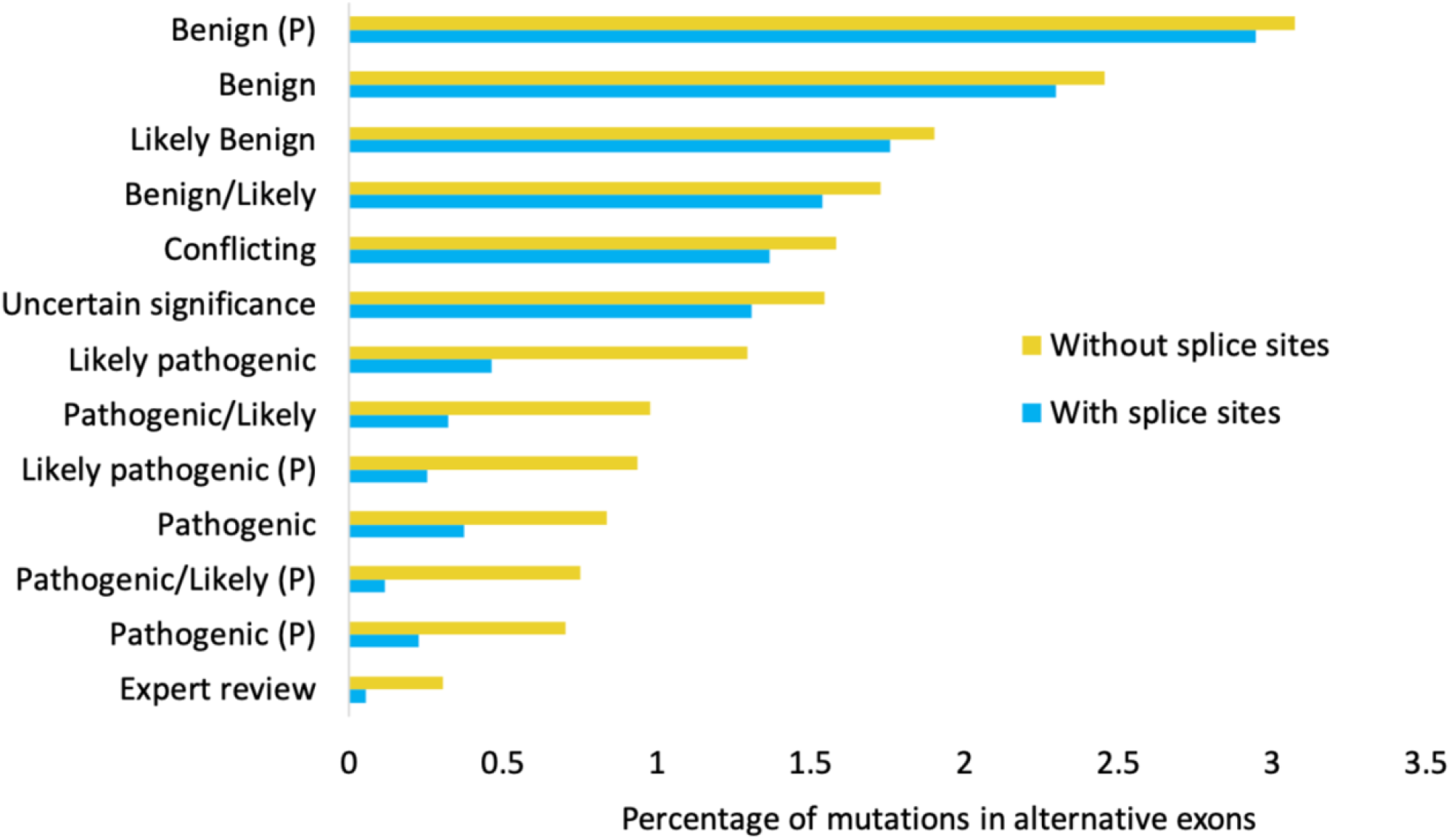
The percentage of ClinVar mutations that map to alternative CDS. The percentage of mutations that map to alternative rather than APPRIS principal CDS by groups. Mutations grouped by ClinVar labels; the labels correspond to the CLIN_SIG entry. Labels with a “P” are also supported by PubMed entries according to ClinVar. “Expert review” are those mutations labelled as “reviewed by expert panel”. Yellow bars show the percentage of mutations in alternative isoforms when mutations are mapped strictly to CDS boundaries, blue bars when mutations are mapped to to CDS boundaries plus those nucleotides likely to form part of the conserved intronic splice site.

Restricting mutations to those that are supported by PubMed references seems to magnify differences between pathogenic and benign mutations. The proportion of pathogenic and likely pathogenic mutations mapping to alternative exons decreases (from 0.89% to 0.73% for mutations tagged as “Pathogenic”, and from 1.35% to 0.94% for mutations tagged as “Likely pathogenic”), while the proportion of benign mutations that map to APPRIS alternative exons increases (2.51% to 3.39%) among variants with PubMed references,

Preliminary analysis of the results suggested that many of the mutations that mapped to alternative coding regions did so in exons generated from 5’ and 3’ alternative splice sites and intron retentions and that many of these were within two or three nucleotides of the splice site of the principal CDS. It is highly probable that the phenotypic effect of these mutations is caused by interference with the splicing mechanism of the principal coding exon rather than by the change of an amino acid in the alternative isoform [ref?]. So, we carried out a second, intronic splice site motif aware mapping of ClinVar mutations to coding exons. We extended coding exons in principal transcripts by three nucleotides at the 5’ end of the exon and 5 nucleotides at the 3’ end. We counted the mutations that mapped to these extended CDS as mutations to APPRIS principal transcripts instead of to the alternative transcripts as we had in the first analysis.

The effect was a substantial decrease in the proportion of mutations that mapped to alternative coding exons. Another 1,804 more ClinVar mutations were captured by principal exons with this splice-site motif aware mapping, with just 1.32% of all variants mapping to alternative exons. The reduction was not uniform across all classes of mutations (Figure 4). Benign mutations that mapped to alternative CDS fell by less than 10% (by 6.8% across all Benign mutations, by just 3.7% among those Benign mutations with PubMed support), but pathogenic mutations that mapped to alternative CDS were cut by more than half. Splice site motif aware mapping cut numbers of all pathogenic mutations mapping to alternative coding exons by 52.9%, while the number of pathogenic mutations with PubMed support that mapped to alternative coding exons fell by 65.1%. The largest reduction was in those mutations reviewed by an expert panel where the numbers of mutations that mapped to alternative isoforms were slashed by 82.8%; just 5 of 9,491 expert reviewed mutations map to alternative CDS.

The American College of Medical Genetics and Genomics (ACMG) recommend that laboratories should not limit themselves to the longest transcript, but should also evaluate the impact of mutations on alternate transcripts or extended untranslated regions [8], though at the same time, they warn against over-interpreting truncating variants that fall in alternative exons. Researchers are reminded to be cautious when assigning clinical significance to mutations in exons where no other pathogenic variants exist [8].

Referring mutations to a reference transcript, and being cautious about variant interpretation in novel exons, do make it slightly less likely that mutations are predicted as pathogenic in alternative transcripts. Nevertheless, the advice is clear - if a mutation in an alternative exon has a pathogenic effect, it should be considered to be pathogenic.

### Manual curation of pathogenic mutations in APPRIS alternative exons

Pathogenic mutations supported by PubMed publications map almost exclusively to APPRIS principal variants rather than alternative transcripts. There were 34,833 mutations classified as Pathogenic, Likely pathogenic or Pathogenic/likely pathogenic by ClinVar with PubMed support. After taking into account mutations that mapped to principal exon splice sites, just 76 mapped to alternative exons (0.22%). These mutations are listed in Supplementary table 1.

We carried out a manual review of these 76 pathogenic mutations, using the supporting publications to determine whether or not the effect of the mutation was on coding DNA, and whether or not the mutation was correctly traduced between research paper and clinical database. We found that just 48 of the 76 mutations appeared to manifest their pathogenicity through a mutation of the functional alternative protein. That is, in just 48 of these 76 mutations the predicted pathogenic effect was likely to result in a change to the sequence – and therefore the function – of an alternative protein isoform. These 48 validated mutations map to 30 distinct alternative transcripts and make up fewer than 0.14% of all 34,833 PubMed-supported pathogenic mutations.

Of the 28 pathogenic mutations that we rejected there were 7 reported mutations for which we could find no supporting evidence in the cited PubMed papers. It is possible that some of these mutations are pathogenic mutations and there are better supporting references that do describe the position of the mutations. These predicted pathogenic mutations were in alternative exons in genes *GPR161, KARS1, LTBP4, MEF2C, NEK1, RPGRIP1L*, and *VPS13B*.

Published papers for another 9 of the pathogenic mutations in alternative exons show that the mutations affect non-coding features rather than alternative coding exons. For example, 3 of the mutations are in a nonsense-mediated decay (NMD) poison exon in *SNRPB* [47]. The mutation in *HBD* falls in a GATA1 binding site that also coincides with an alternative coding exon [48]. The mutation in *SDCCAG8* is in an exonic splicing enhancer [49] that overlaps an annotated alternative coding exon.

In all these cases the mutation is likely to be pathogenic because of its effect on a non-coding feature, but database curators have annotated coding exons overlapping these important non-coding regions. These coding exons are only conserved in primates and generally have little transcript support. Apart from the poison exon in *SNRPB*, these coding exons are good candidates to be eliminated from the human reference set.

Two mutations affect the splicing of the main transcript. The mutation in *CYP21A2* introduces a novel NMD-inducing splice site rather than affecting the alternative annotated coding exon that is annotated in the same position [50]. Aberrant splicing is well studied [51] and often has a pathogenic effect. Aberrant splicing can result from both mutations to constitutive splice sites and mutations that generate novel splice sites. The mutation in *DNA2* maps to an alternative coding exon, but also falls 7 nucleotides downstream of exon 20 of the main variant. Most intronic splice sites are conserved by no more than 5 nucleotides, but exon 20 in *DNA2* has a U12-type splice site. U12-type splice sites make up fewer than 0.5% of 5’ introns. They also have 7 conserved nucleotides [52], so they would not be caught by our splice site motif aware mapping. The authors of the paper on the *DNA2* mutation [53] confirm that the mutation leads to the skipping of exon 20 in the principal transcript.

Three annotations appear to be erroneous. For example, the authors of the paper on a mutation in the alternative coding exon in *KCNQ2* show that the mutation is in the coding exon of the principal variant [54], but in ClinVar the mutation is mis-annotated at an equivalent residue position in the alternative exon.

Finally, seven of the pathogenic mutations predicted to have their effect on alternative isoforms are not supported by cross-species conservation or transcript evidence. Almost all alternative exons with validated pathogenic mutations are highly conserved - all but one of the 48 validated pathogenic mutations are in alternative exons that are conserved at least across eutherian species - while two thirds of the predicted pathogenic variants where the published evidence does not support a pathogenic effect on the alternative isoform are in primate-derived exons, most of these conserved only across higher primates. Recently evolved alternative exons lacking experimental evidence of expression are unlikely to have gained sufficient functional importance for any mutations to have a pathogenic effect [22].

One example of a pathogenic mutation in a primate-derived alternative exon is in the gene *ACTG2* which produces smooth muscle actin. The main isoform of *ACTG2* is highly conserved. Apart from the first three residues, the sequence is practically identical across all vertebrate species and is even 95% identical to muscle actin in invertebrates. The predicted pathogenic mutation affects alternative transcript ENST00000409918, which is forecast to produce a truncated protein that substitutes almost three quarters of the known protein structure, and more than three quarters of the ATP binding residues [Figure 5], for a new 34 residue C-terminal tail. This novel 3’ CDS would theoretically produce a protein isoform with a function that will not even slightly resemble that of actin. This alternative transcript is annotated in the “Basic” quality set in GENCODE, but the novel C-terminal is derived from a LINE2 transposon and is conserved only in chimpanzee.

**Figure 5.**
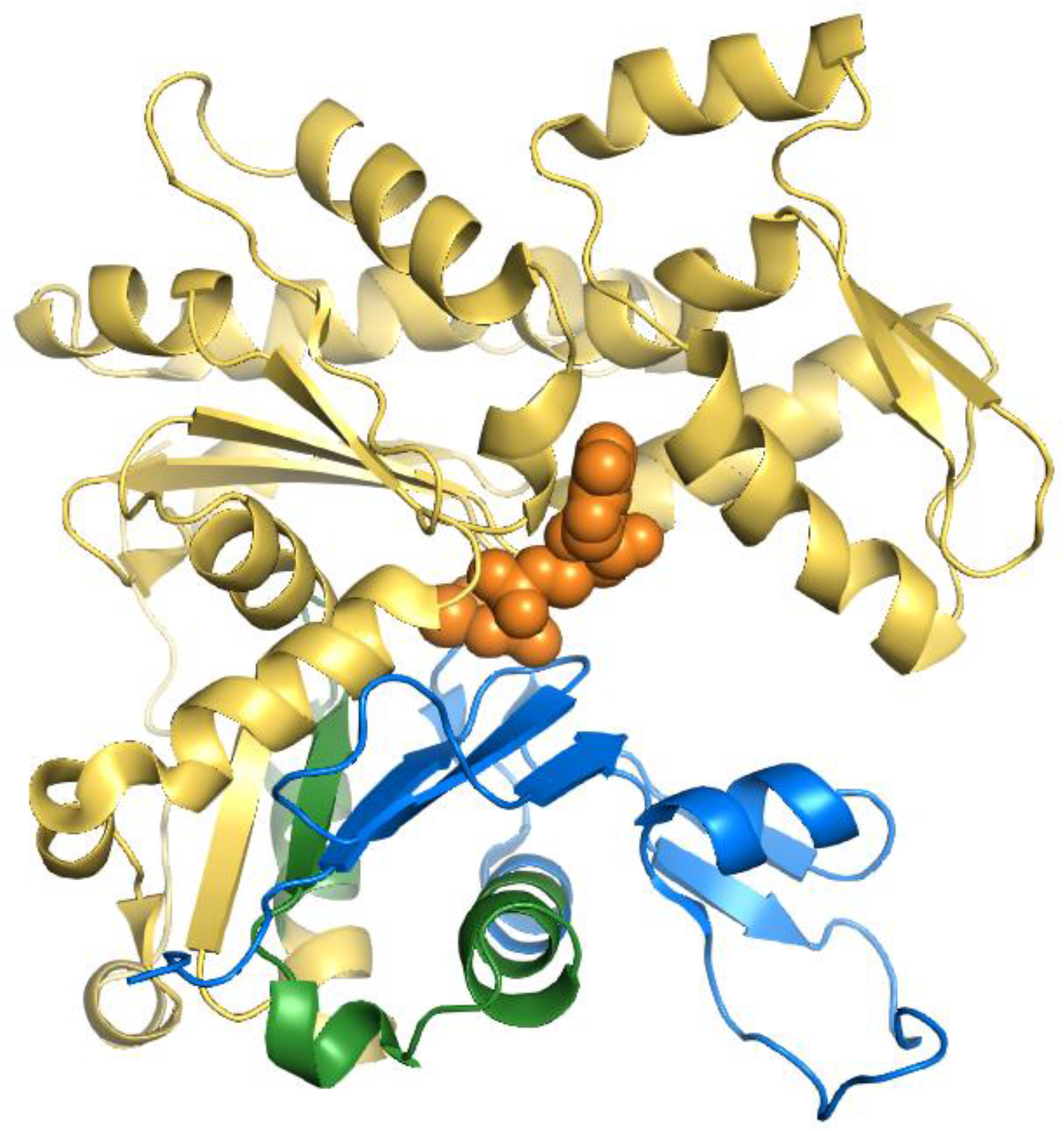
The *ACTG2* alternative variant mapped onto the structure of smooth muscle actin. The sequence of alternative isoform ENSP00000387182 mapped onto the structure of chicken smooth muscle actin (PDB: 3w3d). The region that is maintained in the alternative isoform is shown in blue, while the 34 residues that would be replaced by the novel primate exon with the pathogenic mutation are shown in green. The remainder of the structure (in yellow) would be lost from this presumed protein. The ATP and calcium bound by the chicken actin protein (almost certainly lost in the alternative isoform) are in orange space fill. Mapping was carried out using HHPRED and the image generated from PyMol.

It is hard to imagine how a mutation to this truncated, transposon-derived protein isoform could play an important role in megacystis-microcolon-intestinal hypoperistalsis syndrome, a severe disorder that affects bladder and intestine muscles. The transcript also appears not to be expressed in any tissue [55] and certainly not in bladder or intestines. The authors of the associated paper [56] do not produce any supporting evidence apart from association studies, but they do admit that “the data from this family suggests but perhaps do not prove entirely that the alternative exon 4 which would result in a very short protein isoform is functionally important”.

Another example is the mutation in *SCN1B*. It is located in an exon with no conservation that is partly based on a SINE Alu transposon and that appears to have no expression at all [55]. It is hard to square this evidence with a supposed role in heart regulation [57]. The mutation in the alternative isoform of *LDB3* would change isoleucine to methionine in a little-used alternative exon. Half of the exon has little conservation beyond primates and the isoleucine itself is not conserved beyond primates. What little expression the exon has is concentrated in testis, and there is no evidence that it is expressed in heart [55], which is incongruent given that the mutation is supposed to be a cause of dilated cardiomyopathy [58].

The pathogenic mutation in *NOBOX* is presumed pathogenic because, like the other pathogenic mutants in the paper [59], it falls within the boundaries of a conserved homeobox domain. However, in this case the serine to threonine mutation is inside a primate-derived alternative exon that itself breaks the conserved homeobox domain in two. This inserted exon is adjacent to the conserved asparagine and arginine residues that bind the DNA (Figure 6), so the inserted exon itself is likely to eliminate DNA binding. This alternative exon also has little or no evidence for expression [55].

**Figure 6.**
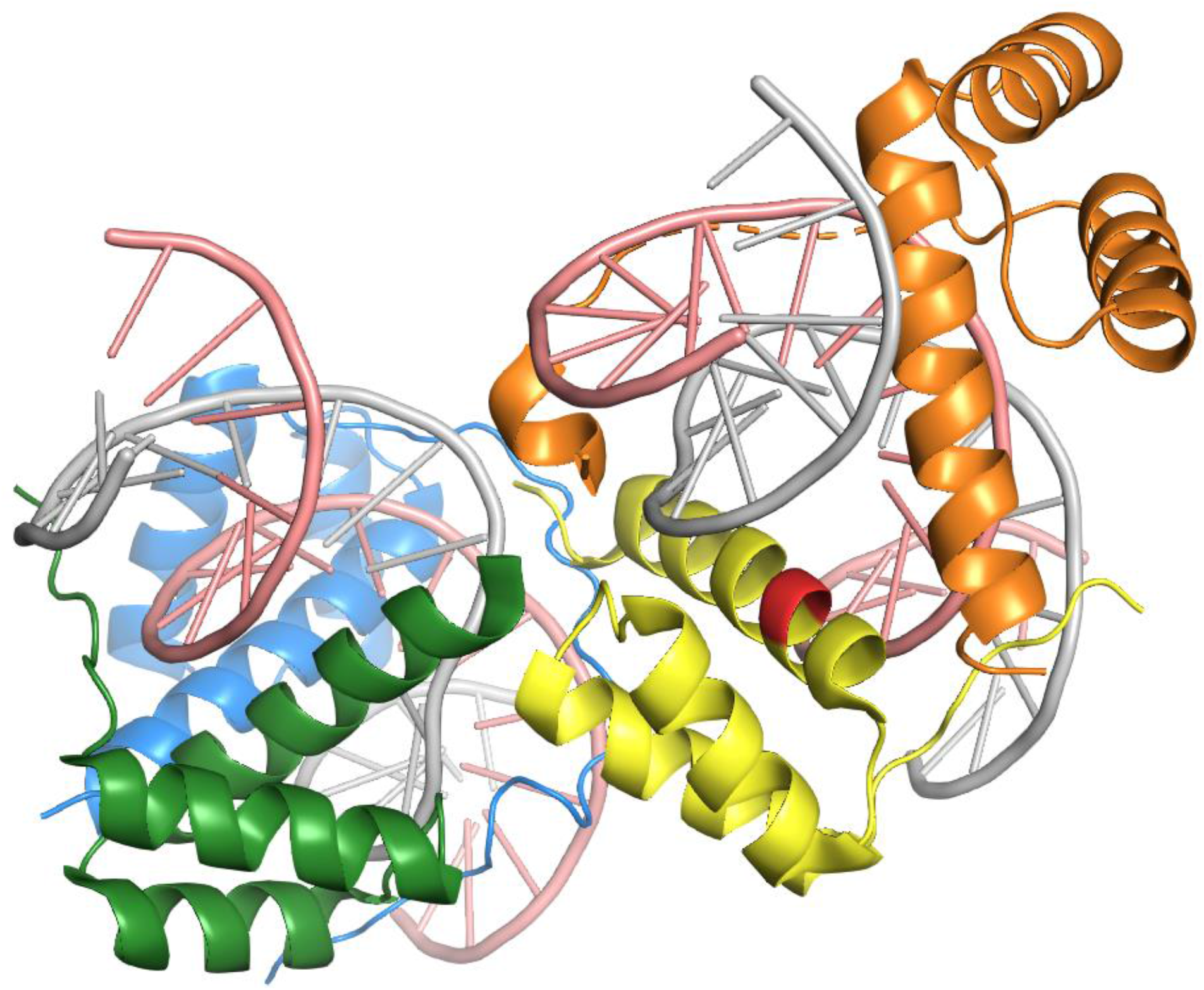
The effect of the inserted exon in the *NOBOX* alternative isoform. The image shows the crystal structure of *Drosophila melanogaster Aristaless* and *Clawless* homeodomain proteins bound to DNA (PDB:3A01). The *Aristaless* protein (chain F, yellow) has 57% identity to the *NOBOX* homeobox domain. The residues where the 32 amino acid insert would break the *NOBOX* homeobox domain are highlighted in red. The insertion is right next to the conserved DNA-binding residues of the homeobox domain and this primate derived exon would almost certainly banish the *NOBOX* homeobox DNA-binding function. Mapping was carried out using HHPRED and the image generated from PyMol.

Pathogenic mutations in some recently evolved alternative exons do appear to affect the alternative protein. The alternative transcript in *ERCC6* evolved in the primate lineage, though the alternative exon was co-opted from a transposon with a last common ancestor in fish [32]. The alternative exon that houses the pathogenic mutation in *REEP6* evolved during the eutherian clade. However, it is quite clearly brain-specific [55] and is part of the main transcript variant in retina [4]. The mutation is predicted to cause autosomal-recessive retinitis pigmentosa [60].

### Pathogenic mutations in exons alternative to MANE Select and longest variants

We carried out the same analysis with the GENCODE v37 MANE Select transcripts and the longest transcripts for each gene. MANE Select transcripts do not yet cover all coding genes, but genes with MANE Select transcript do cover more than 95% of the pathogenic mutations in ClinVar. In genes with a defined MANE Select transcript, MANE Select transcripts captured all but 67 of the 33,736 pathogenic mutations with PubMed support. Just 0.2% of ClinVar pathogenic mutations with PubMed support map to transcripts considered by MANE to be alternative. This result was highly similar to what we found in APPRIS, and indeed many of the mutations overlapped in the two sets.

There was no significant difference between the proportion of pathogenic variants captured by APPRIS principal splice variants and those captured by MANE Select transcripts. However, analysis of the longest isoform of each coding gene did find significantly more pathogenic and likely pathogenic mutations mapping to alternative shorter isoforms. A total of 160 of the 34,833 ClinVar pathogenic or likely pathogenic mutations supported by PubMed publications (0.46%) were not captured by the longest isoform (Fisher exact test, p < 0.00001).

We also carried out a manual curation of these pathogenic mutations. We validated 47 of the 67 pathogenic mutations affecting MANE Select alternative exons, again similar to what we found using APPRIS to select the main variants. We validated 144 of 160 pathogenic mutations that mapped to those (shorter) isoforms that were alternative to the longest isoform. Three times as many validated pathogenic mutations map to alternative exons when the longest isoform is used to determine the most important isoform instead of the MANE Select or APPRIS principal splice variants, despite the fact that the longest variants include more coding sequence. Pathogenic mutations in alternative exons from these two studies can be found in the Supplementary materials.

### Combining APPRIS principal and MANE Select variants

We looked at the combined power of APPRIS principal and MANE Select transcripts. For those genes where both methods agreed on the main splice variant, 33,631 of 33,670 pathogenic mutations with PubMed support (99.9%) mapped to exons from APPRIS principal and MANE Select transcripts. Just 39 pathogenic mutations mapped to exons from transcripts designated as alternative both by APPRIS and the expert RefSeq and GENCODE curators (0.11%). Just 22 of these pathogenic mutations (0.065%) are validated by their PubMed references. These 22 pathogenic mutations map to exons in just 15 GENCODE v37 alternative transcripts. When they agree, APPRIS principal variants and MANE Select transcripts capture almost all annotated clinically important mutations.

### Few alternative transcripts have pathogenic mutations in ClinVar

In order to quantify the number of alternative transcripts that have ClinVar pathogenic and likely pathogenic mutations, we extended our analysis to include pathogenic mutations regardless of whether or not they were supported by a publication. To guarantee that at least one of the pathogenic mutations maps to an alternative isoform, we counted the number of genes in which pathogenic mutations mapped to more than one coding sequence.

After we took into account mutations that affect intronic splice site motifs of principal exons, just 54 genes had ClinVar pathogenic or likely pathogenic mutations in two or more distinct coding transcripts. Many of these predicted pathogenic mutations are without PubMed support, so are impossible to analyse manually. However, we were able to evaluate the relative age of the exons with pathogenic mutations and determine their experimental support. As we have already shown, recently evolved alternative exons with little experimental support are unlikely candidates for pathogenic mutations that affect alternative proiteins.

Within the 54 genes there were pathogenic mutations in 59 alternative exons. The mutations mapped to three or more exons in distinct coding sequences in four genes, *GCNT2, SCN1A, PCDH15* and *GNAS*. We found that 27 of the 59 exons were little expressed, primate-derived or both. Five of these exons would lead to NMD, including all three alternative exons in *SCN1A*, and 11 derived from primate transposons. Eleven of these 27 alternative exons were generated by alternative 3’ or 5’ splice sites. This could be the result of the ACMGC maxim that encourages the annotation of pathogenic mutations in exons that already have pathogenic mutations [8], since all these splice events elongate the principal exon.

This left us with just 32 examples where both principal and alternative transcripts were annotated with pathogenic mutations. The 32 alternative transcripts are annotated in 29 genes because there are pathogenic mutations in three distinct coding transcripts in *GCNT2, GNAS* and *PCDH15*. We classed these 32 alternative transcripts with pathogenic mutations as “validated”. Even though we were not able to validate all the annotations via PubMed publications, 12 of the 32 pathogenic mutations overlapped with the previous sets, so were supported by experiments.

### Distinguishing functionally important alternative isoforms

It is abundantly clear that APPRIS principal and/or MANE Select variants best represent the main functional variant in the cell. They should be regarded as the main coding variant for each gene when annotating clinical variants. But alternative variants can also be clinically important. How can researchers distinguish clinically important alternative isoforms from those protein isoforms that are unlikely to be affected by pathogenic mutations?

There are cues within the 32 validated alternative transcripts with pathogenic mutations. More than half of the 32 splice events that generate the alternative transcripts involve the alternative splicing of tandem duplicated exons [**3, 61**], even though tandem duplicated exon substitutions make up only a tiny fraction of annotated splice events [**3**]. Four splice variants are generated by swapping one type of Pfam domain for another, and these splice events are even more rare in the human genome. Most importantly, two thirds of the 32 examples of pathogenic mutations in alternative exons have support from both cross-species conservation and expression; more than 90% of the alternative exons with validated pathogenic mutations can be traced back prior to the evolution of mammals.

For example, the mutation in the alternative exon in *SLC25A3* is predicted to cause mitochondrial phosphate carrier deficiency, a fatal defect characterised by lactic acidosis, hypertrophic cardiomyopathy, and muscular hypotonia [**62**]. This exon is not expressed outside or heart or muscle, but crucially it is part of the main variant in muscle and heart [**4, 55**] and the exon is conserved in lamprey [**3**], so its evolution pre-dated the earliest vertebrates. ClinVar records a pathogenic mutation in the alternative tandem duplicated exon of *TCF3* that is associated with agammaglobulinemia [**63**]. The tandem duplication that generated this event occurred more than 560 million years ago and both exons have widespread expression, suggesting that both principal and alternative isoforms are functionally important.

Ensembl/GENCODE and RefSeq manual annotators have started to annotate clinically important alternative variants based on the annotations in ClinVar. Alternative transcripts in *SLC25A3* and *TCF3* are annotated as MANE Plus Clinical transcripts, as are another 20 of the 32 alternative transcripts with validated pathogenic mutations. Currently there are 64 MANE Plus Clinical transcripts with known clinically important variants.

We developed TRIFID to predict likely functionally important alternative isoforms [**32**]. In order to determine the usefulness of the TRIFID score in predicting the clinical importance of alternative exons, we evaluated the TRIFID scores for alternative transcripts with pathogenic mutations. We combined the alternative transcripts from the 32 pairs of principal and alternative transcripts with validated pathogenic mutations and the 30 APPRIS alternative transcripts that had validated pathogenic mutations with PubMed. In total, there were 51 alternative transcripts because 11 appeared in both lists. Just over half (26) had a normalised TRIFID score of over 0.8, while just two had TRIFID scores of below 0.2.

As a control, we calculated the best TRIFID score for all alternative exons defined by APPRIS as alternative. For each alternative exon, we took the best scoring coding variant that it was annotated in. We binned all annotated alternative exons in the human reference set by their normalised TRIFID scores. Almost 85% of the 76,134 alternative coding exons in GENCODE v37 had a TRIFID score of less than 0.2, while just 3.3% scored more than 0.8.

The proportion of annotated alternative coding exons with validated pathogenic mutations in each bin are shown in **Figure 7**. The higher the TRIFID score, the more likely an alternative variant was to harbour a pathogenic mutation affecting protein function. Just 0.003% of alternative exons from transcripts with TRIFID scores below 0.2 had validated pathogenic mutations. By contrast, validated pathogenic mutations were found in more than 1% of those alternative exons that came from transcripts with TRIFID scores between 0.8 and 1. Remarkably, exons from the highest scoring TRIFID alternative transcripts have 332 times as many validated pathogenic mutations as the 85% of exons in the lowest scoring bin.

**Figure 7.**
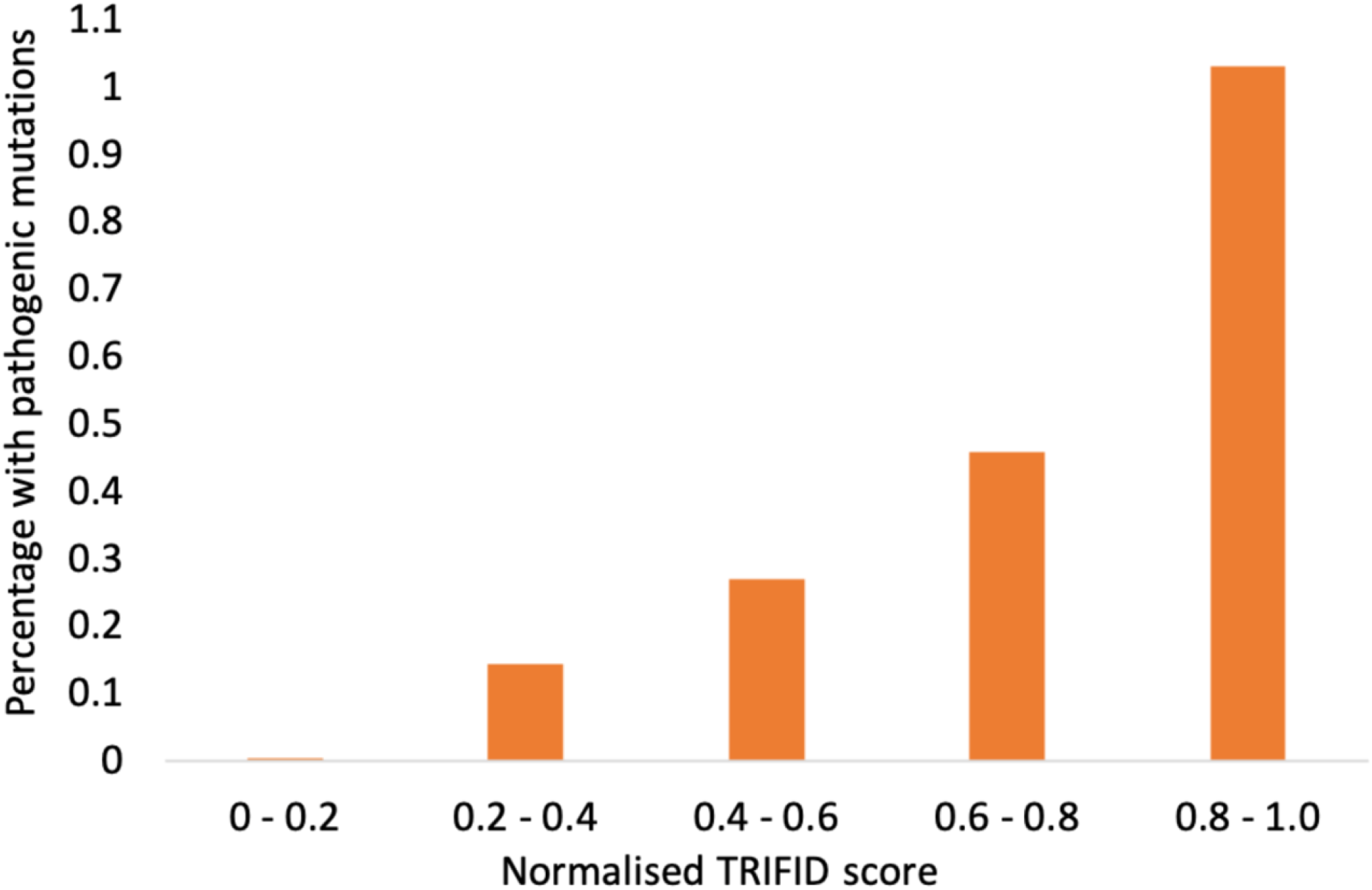
Percentage of alternative exons with pathogenic mutations binned by TRIFID score. We validated 51 pathogenic mutations in exons determined by APPRIS to be in alternative variants. We binned all alternative coding exons in the human reference set by the TRIFID score of the best-scoring transcript in which they were annotated, and calculated the percentage of exons with validated pathogenic mutations in each bin. Despite the low numbers of validated pathogenic mutations, Fisher exact tests showed that the percentage of pathogenic mutations with best TRIFID scores exceeding 0.8 was significantly higher than those in all other bins, and that the percentage of pathogenic mutations among exons with TRIFID scores below 0.2 was significantly lower than all other bins.

There are pathogenic mutations in two transcripts that score less than 0.2 in TRIFID. The mutation in *GNAS* is in the huge 5’ CDS. Much of the exon is conserved in fish, but other parts of the exon are not conserved and produce a region of low complexity. Since TRIFID considers splice isoforms as a whole, this alternative isoform scores poorly because of this non-conserved region.

The other alternative variant (in *BNC2*) is interesting because it is part of an unfinished gene model in GENCODE v37. At the moment the pathogenic mutation falls in an alternative 3’ CDS that is also missing a 5’ CDS. The CDS in which the exon falls evolved during the mammalian clade and is almost 100% conserved across mammalian species. In all these species the annotated exon is 66-bases long. In humans the exon is just 62 bases long and the change of reading frame introduces a premature stop codon and leads to the loss of the Zf-C2H2 domain. As a result, TRIFID gives this variant a poor score. The mammalian splice site is conserved in humans, so when the 66-base, frame maintaining exon is annotated as part of a full-length transcript, the transcript will get a much better TRIFID score.

## Conclusions

We find that peptide evidence and clinical variants support the hypothesis that most coding genes have single main protein isoform. This main cellular isoform is best described by the MANE Select transcripts and the APPRIS principal isoforms. Peptide evidence from proteomics experiments is the best available evidence for the translation into protein of coding transcripts, and APPRIS principal isoforms and MANE Select variants each agree with the main proteomics isoform over approximately 95% of the almost 7,000 genes for which we were able to determine a main isoform. Where MANE Select and APPRIS P1 principal isoforms coincide (5,888 genes), the agreement with the main proteomics isoform is 98.2%. Main isoforms from proteomics experiments, APPRIS principal isoforms and MANE Select transcripts are three orthogonal means of determining a main splice variant and the fact that they agree over such a high proportion of genes reaffirms that most coding genes have a single main cellular isoform [2].

Data from the ClinVar database shows that pathogenic mutations map almost exclusively to a single transcript variant per gene. The transcripts almost always coincide with the APPRIS principal isoforms and the MANE Select transcripts. The number of pathogenic mutations in ClinVar that map to alternative exons is much lower than would be expected. We also found that the vast majority of the pathogenic mutations that initially mapped to alternative exons either affected the splicing of the main transcript [51], affected conserved non-coding features, or were not supported by cross-species conservation or experimental data. In fact, just 0.065% of all pathogenic mutations supported by PubMed publications affected protein isoforms that APPRIS and MANE Select tag as alternative. This is further evidence to support the theory that most protein coding genes have a single main protein isoform.

While most coding genes have one main isoform, some genes do have multiple functionally important splice variants. This is also clear from the proteomics experiments here and in previous analyses [3, 4, 30], and it is important to be able to identify these functionally important isoforms. We show that two related tools capture clinical importance in alternative variants. MANE Plus Clinical transcripts are curated by Ensembl/GENCODE and RefSeq annotators and capture most pathogenic mutations that are already annotated in alternative exons. TRIFID, which forms part of the APPRIS database, predicts the functional importance of protein isoforms.

We have already shown that higher scoring TRIFID isoforms are more likely to be functionally important [32]. These results show that transcript variants with a higher associated TRIFID score are also much more likely to be clinically relevant. Alternative exons in high scoring TRIFID transcripts are more than 300 times more likely than those from low scoring transcripts.to harbour pathogenic mutations. APPRIS, TRIFID and MANE are available for both the Ensembl/GENCODE and RefSeq annotations of the human gene set. APPRIS principal isoforms and TRIFID functional importance scores are also available for further vertebrate and invertebrate model species.

We believe that our work shows that the researchers should be more aware of the correct main isoform when making predictions for pathogenicity. Mutations that fall in alternative variants are already treated with care [8], but researchers ought to be aware of the likely main isoform of each coding gene. The longest isoform is not a substitute for the main isoform and is a particular problem in the RefSeq gene set. Many RefSeq transcripts are computational predictions and a lot of these are without supporting evidence. The longest transcript in many RefSeq coding genes will be a computational prediction. We have shown that both MANE Select transcripts and APPRIS principal isoforms are good predictors of the main cellular isoform and are particularly powerful when they agree. MANE Select transcripts and APPRIS principal isoforms should be incorporated into clinical variant interpretation.

## Acknowledgements

We would like to thank Thomas A. Walsh for proteomics analysis work carried out.

## References

1. Frankish, A., Diekhans, M., Ferreira, A.M., Johnson, R., Jungreis, I., Loveland, J., Mudge, J.M., Sisu, C., Wright, J., Armstrong, J. et al. (2019) GENCODE reference annotation for the human and mouse genomes. Nucleic Acids Res., 47, D766–D773.

2. Ezkurdia, I., Rodriguez, J.M., Carrillo-de Santa Pau, E., Vázquez, J., Valencia, A. and Tress, M.L. (2015) Most highly expressed protein-coding genes have a single dominant isoform. J. Proteome Res., 14, 1880–1887.

3. Martinez Gomez L, Pozo F, Walsh TA, Abascal F, Tress ML. The clinical importance of tandem exon duplication-derived substitutions. Nucleic Acids Res. 2021 49:8232–46.

4. Rodriguez, J.M., Pozo, F., di Domenico, T., Vazquez, J. and Tress, M.L. (2020) An analysis of tissue-specific alternative splicing at the protein level. PLoS Comp. Biol., 16, e1008287.

5. Tress, M.L., Abascal, F. and Valencia, A. (2017) Alternative Splicing May Not Be the Key to Proteome Complexity. Trends Biochem. Sci., 42, 98–110.

6. Tress, M.L., Abascal, F. and Valencia, A. (2017) Most alternative isoforms are not functionally important. Trends Biochem Sci., 42, 408–410.

7. Richards CS, Bale S, Bellissimo DB, Das S, Grody WW, Hegde MR, Lyon E, Ward BE; Molecular Subcommittee of the ACMG Laboratory Quality Assurance Committee. ACMG recommendations for standards for interpretation and reporting of sequence variations: Revisions 2007. Genet Med. 2008 10:294–300.

8. Richards S, Aziz N, Bale S, Bick D, Das S, Gastier-Foster J, Grody WW, Hegde M, Lyon E, Spector E, Voelkerding K, Rehm HL; ACMG Laboratory Quality Assurance Committee. Standards and guidelines for the interpretation of sequence variants: a joint consensus recommendation of the American College of Medical Genetics and Genomics and the Association for Molecular Pathology. Genet Med. 2015 17:405–24.

9. Sayers, E.W., Beck, J., Brister, J.R., Bolton, E.E., Canese, K., Comeau, D.C., Funk, K., Ketter, A., Kim, S., Kimchi, A. et al. (2020) Database resources of the National Center for Biotechnology Information. Nucleic Acids Res., 48, D9–D16.

10. Bione S, D’Adamo P, Maestrini E, Gedeon AK, Bolhuis PA, Toniolo D. A novel X-linked gene, G4.5. is responsible for Barth syndrome. Nat Genet. 1996 12:385–9.

11. Xu Y, Zhang S, Malhotra A, Edelman-Novemsky I, Ma J, Kruppa A, Cernicica C, Blais S, Neubert TA, Ren M, Schlame M. Characterization of tafazzin splice variants from humans and fruit flies. J Biol Chem. 2009 284:29230–9.

12. Schlame M, Xu Y. The Function of Tafazzin, a Mitochondrial Phospholipid-Lysophospholipid Acyltransferase. J Mol Biol. 2020 432:5043–5051.

13. Barth PG, Scholte HR, Berden JA, Van der Klei-Van Moorsel JM, Luyt-Houwen IE, Van ‘t Veer-Korthof ET, Van der Harten JJ, Sobotka-Plojhar MA. An X-linked mitochondrial disease affecting cardiac muscle, skeletal muscle and neutrophil leucocytes. J Neurol Sci. 1983 62:327–55.

14. The UniProt Consortium. (2017) UniProt: the universal protein knowledgebase Nucleic Acids Res., 45, D158–D159.

15. Vaz FM, Houtkooper RH, Valianpour F, Barth PG, Wanders RJ. Only one splice variant of the human TAZ gene encodes a functional protein with a role in cardiolipin metabolism. J Biol Chem. 2003 278:43089–94.

16. MacArthur JA, Morales J, Tully RE, Astashyn A, Gil L, Bruford EA, Larsson P, Flicek P, Dalgleish R, Maglott DR, Cunningham F. Locus Reference Genomic: reference sequences for the reporting of clinically relevant sequence variants. Nucleic Acids Res. 2014 42:D873–8.

17. Finn, R.D., Coggill, P., Eberhardt, R.Y., Eddy, S.R., Mistry, J., Mitchell, A.L., Potter, S.C., Punta, M., Qureshi, M., Sangrador-Vegas, A., et al. (2016) The Pfam protein families database: towards a more sustainable future. Nucleic Acids Res., 44, D279–D285.

18. Landrum, M.J., Lee, J.M., Benson, M., Brown, G.R., Chao, C., Chitipiralla, S., Gu, B., Hart, J., Hoffman, D., Jang, W. et al. (2018) ClinVar: improving access to variant interpretations and supporting evidence. Nucleic Acids Res., 46, D1062–D1067.

19. Tunyasuvunakool K, Adler J, Wu Z, Green T, Zielinski M, Žídek A, Bridgland A, Cowie A, Meyer C, Laydon A, et al. Highly accurate protein structure prediction for the human proteome. Nature. 2021 Jul 22.

20. Lin, M.F., Jungreis, I. and Kellis, M. (2011) PhyloCSF: A comparative genomics method to distinguish protein coding and non-coding regions. Bioinformatics, 27, 275–282.

21. Pollard KS, Hubisz MJ, Rosenbloom KR, Siepel A. Detection of nonneutral substitution rates on mammalian phylogenies. Genome Res. 2010 Jan;20(1):110–21.

22. Martinez-Gomez, L., Abascal, F., Jungreis, I., Pozo, F,. Kellis, M., Mudge, J.M. and Tress, M.L. (2020) Few SINEs of life: Alu elements have little evidence for biological relevance despite elevated translation. NAR Genom. Bioinform., 2, lqz023.

23. Hijikata A, Yura K, Ohara O, Go M. Structural and functional analyses of Barth syndrome-causing mutations and alternative splicing in the tafazzin acyltransferase domain. Meta Gene. 2015 4:92–106.

24. Gonzàlez-Porta, M., Frankish, A., Rung, J., Harrow, J. and Brazma, A. (2013) Transcriptome analysis of human tissues and cell lines reveals one dominant transcript per gene. Genome Biol., 14, R70.

25. Li, H.D., Menon, R., Govindarajoo, B., Panwar, B., Zhang, Y., Omenn, G.S. and Guan, Y. (2015) Functional Networks of Highest-Connected Splice Isoforms: From The Chromosome 17 Human Proteome Project. J. Proteome Res., 14, 3484–3491.

26. Carlyle BC, Kitchen RR, Zhang J, Wilson RS, Lam TT, Rozowsky JS, Williams KR, Sestan N, Gerstein MB, Nairn AC. Isoform-Level Interpretation of High-Throughput Proteomics Data Enabled by Deep Integration with RNA-seq. J Proteome Res. 2018 17:3431–3444.

27. Harte, R.A., Farrell, C.M., Loveland, J.E., Suner, M.M., Wilming, L., Aken, B., Barrell, D., Frankish, A., Wallin, C., Searle, S, et al. (2012) Tracking and coordinating an international curation effort for the CCDS Project. Database, 2012, 1–12.

28. Rodriguez, J.M., Rodriguez-Rivas, J,. Di Domenico, T., Vázquez, J., Valencia, A. and Tress, M.L. (2018) APPRIS 2017: Principal isoforms for multiple gene sets. Nucleic Acids Res., 46, D213–D217.

29. Howe, K. L., Achuthan, P., Allen, J., Allen, J., Alvarez-Jarreta, J., Amode, M. R., Armean, I. M., Azov, A. G., Bennett, R., Bhai, J., Billis, K., Boddu, S., Charkhchi, M., Cummins, C., Da Rin Fioretto, L., Davidson, C., Dodiya, K., El Houdaigui, B., Fatima, R., Gall, A., … Flicek, P. (2021). Ensembl 2021. Nucleic acids research, 49, D884–D891.

30. Abascal, F., Ezkurdia, .I, Rodriguez-Rivas, J., Rodriguez, J.M., del Pozo, A., Vázquez, J., Valencia, A. and Tress, M.L. (2015) Alternatively Spliced Homologous Exons Have Ancient Origins and Are Highly Expressed at the Protein LFevel. PLoS Comp. Biol., 11, 1–29.

31. Liu, T. and Lin, K. (2015) The distribution pattern of genetic variation in the transcript isoforms of the alternatively spliced protein-coding genes in the human genome. Mol. Biosyst., 11, 1378–1388.

32. Pozo F, Martinez-Gomez L, Walsh TA, Rodriguez JM, Di Domenico T, Abascal F, Vazquez J, Tress ML. Assessing the functional relevance of splice isoforms. NAR Genom Bioinform. 2021 3:lqab044.

33. Kim, M.S., Pinto, S.M., Getnet, D., Nirujogi, R.S., Manda, S.S., Chaerkady, R., Madugundu, A.K., Kelkar, D.S., Isserlin, R., Jain, S. et al. (2014) A draft map of the human proteome. Nature, 509, 575–581.

34. Carlyle BC, Kitchen RR, Kanyo JE, Voss EZ, Pletikos M, Sousa AMM, Lam TT, Gerstein MB, Sestan N, Nairn AC. A multiregional proteomic survey of the postnatal human brain. Nat Neurosci. 2017 Dec;20(12):1787–1795.

35. Bekker-Jensen DB, Kelstrup CD, Batth TS, Larsen SC, Haldrup C, Bramsen JB, Sørensen KD, Høyer S, Ørntoft TF, Andersen CL, Nielsen ML, Olsen JV. An Optimized Shotgun Strategy for the Rapid Generation of Comprehensive Human Proteomes. Cell Syst. 2017 4:587–599.e4

36. Schiza C, Korbakis D, Jarvi K, Diamandis EP, Drabovich AP. Identification of TEX101-associated Proteins Through Proteomic Measurement of Human Spermatozoa Homozygous for the Missense Variant rs35033974. Mol Cell Proteomics. 2019 18:338–351

37. Wang D, Eraslan B, Wieland T, Hallström B, Hopf T, Zolg DP, Zecha J, Asplund A, Li LH, Meng C, et al. A deep proteome and transcriptome abundance atlas of 29 healthy human tissues. Mol Syst Biol. 2019 15:e8503.

38. Deutsch, E.W., Csordas, A., Sun, Z., Jarnuczak, A., Perez-Riverol, Y., Ternent, T., Campbell, D.S., Bernal-Llinares, M., Okuda, S., Kawano, S. et al. (2017) The ProteomeXchange consortium in 2017: supporting the cultural change in proteomics public data deposition. Nucleic Acids Res., 45, D1100–D1106.

39. Abascal, F., Juan, D., Jungreis, I., Martinez, L., Rigau, M., Rodriguez, J.M., Vazquez, J. and Tress, M.L. (2018) Loose ends: almost one in five human genes still have unresolved coding status. Nucleic Acids Res., 46, 7070–7084.

40. Eng, J.K., Jahan, T.A. and Hoopmann, M.R. (2013) Comet: an open-source MS/MS sequence database search tool. Proteomics, 13, 22–24.

41. The, M., MacCoss, M.J., Noble, W.S. and Käll, L. (2016) Fast and Accurate Protein False Discovery Rates on Large-Scale Proteomics Data Sets with Percolator 3.0. J. Am. Soc. Mass. Spectrom., 27, 1719–1727.

42. Ezkurdia, I., Calvo, E., Del Pozo, A., Vázquez, J., Valencia, A. and Tress, M.L. (2015) The potential clinical impact of the release of two drafts of the human proteome. Expert Rev. Proteomics, 12, 579–593.

43. Burley, S.K., Berman, H.M., Kleywegt, G.J., Markley, J.L., Nakamura, H., and Velankar, S. (2017) Protein Data Bank (PDB): The Single Global Macromolecular Structure Archive. Methods Mol. Biol., 1607, 627–641.

44. Gabler, F., Nam, S.Z., Till, S., Mirdita, M., Steinegger, M., Söding, J., Lupas, A.N. and Alva, V. (2020) Protein Sequence Analysis Using the MPI Bioinformatics Toolkit. Curr. Protoc. Bioinformatics, 72, e108.

45. McLaren, W., Gil, L., Hunt, S.E., Riat, H.S., Ritchie, G.R., Thormann, A., Flicek, P. and Cunningham, F. (2016) The Ensembl Variant Effect Predictor. Genome Biol., 17, 122.

46. Rodriguez, J.M., Maietta, P., Ezkurdia, I., Pietrelli, A., Wesselink, J.J., Lopez, G., Valencia, A. and Tress, M.L. (2013) APPRIS: Annotation of principal and alternative splice isoforms. Nucleic Acids Res., 41, 110–117.

47. Lynch DC, Revil T, Schwartzentruber J, Bhoj EJ, Innes AM, Lamont RE, Lemire EG, Chodirker BN, Taylor JP, Zackai EH, et al. Disrupted auto-regulation of the spliceosomal gene SNRPB causes cerebro-costo-mandibular syndrome. Nat Commun. 2014 5:4483.

48. Matsuda M, Sakamoto N, Fukumaki Y. Delta-thalassemia caused by disruption of the site for an erythroid-specific transcription factor, GATA-1, in the delta-globin gene promoter. Blood. 1992 80:1347–51.

49. Otto EA, Hurd TW, Airik R, Chaki M, Zhou W, Stoetzel C, Patil SB, Levy S, Ghosh AK, Murga-Zamalloa CA, et al. Candidate exome capture identifies mutation of SDCCAG8 as the cause of a retinal-renal ciliopathy. Nat Genet. 2010 42:840–50.

50. Higashi Y, Tanae A, Inoue H, Hiromasa T, Fujii-Kuriyama Y. Aberrant splicing and missense mutations cause steroid 21-hydroxylase [P-450(C21)] deficiency in humans: possible gene conversion products. Proc Natl Acad Sci U S A. 1988 85:7486–90.

51. Park E, Pan Z, Zhang Z, Lin L, Xing Y. The Expanding Landscape of Alternative Splicing Variation in Human Populations. Am J Hum Genet. 2018 102:11–26.

52. Sibley, Christopher & Blazquez, Lorea & Ule, Jernej. (2016). Lessons from non-canonical splicing. Nature Reviews Genetics. 17. 10.1038/nrg.2016.46

53. Shaheen R, Faqeih E, Ansari S, Abdel-Salam G, Al-Hassnan ZN, Al-Shidi T, Alomar R, Sogaty S, Alkuraya FS. Genomic analysis of primordial dwarfism reveals novel disease genes. Genome Res. 2014 24:291–9.

54. Richards MC, Heron SE, Spendlove HE, Scheffer IE, Grinton B, Berkovic SF, Mulley JC, Davy A. Novel mutations in the KCNQ2 gene link epilepsy to a dysfunction of the KCNQ2-calmodulin interaction. J Med Genet. 2004 41:e35.

55. Cummings BB, Karczewski KJ, Kosmicki JA, Seaby EG, Watts NA, Singer-Berk M, Mudge JM, Karjalainen J, Satterstrom FK, O’Donnell-Luria AH, et al. Transcript expression-aware annotation improves rare variant interpretation. Nature. 2020 581:452–458.

56. Wangler MF, Gonzaga-Jauregui C, Gambin T, Penney S, Moss T, Chopra A, Probst FJ, Xia F, Yang Y, Werlin S, et al. Heterozygous de novo and inherited mutations in the smooth muscle actin (ACTG2) gene underlie megacystis-microcolon-intestinal hypoperistalsis syndrome. PLoS Genet. 2014 10:e1004258.

57. Watanabe H, Koopmann TT, Le Scouarnec S, Yang T, Ingram CR, Schott JJ, Demolombe S, Probst V, Anselme F, Escande D, et al. Sodium channel β1 subunit mutations associated with Brugada syndrome and cardiac conduction disease in humans. J Clin Invest. 2008 118:2260–8.

58. Vatta M, Mohapatra B, Jimenez S, Sanchez X, Faulkner G, Perles Z, Sinagra G, Lin JH, Vu TM, Zhou Q, et al. Mutations in Cypher/ZASP in patients with dilated cardiomyopathy and left ventricular non-compaction. J Am Coll Cardiol. 2003 42:2014–27.

59. Bouilly J, Bachelot A, Broutin I, Touraine P, Binart N. Novel NOBOX loss-of-function mutations account for 6.2% of cases in a large primary ovarian insufficiency cohort. Hum Mutat. 2011 32:1108–13.

60. Arno G, Agrawal SA, Eblimit A, Bellingham J, Xu M, Wang F, Chakarova C, Parfitt DA, Lane A, Burgoyne T, et al. Mutations in REEP6 Cause Autosomal-Recessive Retinitis Pigmentosa. Am J Hum Genet. 2016 99:1305–1315.

61. Lam SD, Babu MM, Lees J, Orengo CA. Biological impact of mutually exclusive exon switching. PLoS Comput Biol. 2021 17:e1008708.

62. Mayr JA, Merkel O, Kohlwein SD, Gebhardt BR, Böhles H, Fötschl U, Koch J, Jaksch M, Lochmüller H, Horváth R, Freisinger P, Sperl W. Mitochondrial phosphate-carrier deficiency: a novel disorder of oxidative phosphorylation. Am J Hum Genet. 2007 80:478–84.

63. Boisson B, Wang YD, Bosompem A, Ma CS, Lim A, Kochetkov T, Tangye SG, Casanova JL, Conley ME. A recurrent dominant negative E47 mutation causes agammaglobulinemia and BCR(-) B cells. J Clin Invest. 2013 123:4781–5.

